# Single-cell RNA sequencing reveals cellular and molecular heterogeneity in fibrocartilaginous enthesis formation

**DOI:** 10.1101/2023.02.02.526768

**Authors:** Tao Zhang, Wan Liyang, Xiao Han, Linfeng Wang, Jianzhong Hu, Hongbin Lu

## Abstract

The attachment site of the rotator cuff (RC) is a classic fibrocartilaginous enthesis, which is the junction between bone and tendon with typical characteristics of a fibrocartilage transition zone. Enthesis development has historically been studied with lineage tracing of individual genes selected a priori, which does not allow for the determination of single-cell landscapes yielding mature cell types and tissues. Here, we applied Single-cell RNA sequencing (scRNA-seq) to delineate the comprehensive postnatal RC enthesis growth and the temporal atlas from as early as embryonic day 15 up to postnatal week 4. In summary, we compared the development pattern between enthesis and tendon or articular cartilage, then deciphered the cellular heterogeneity and the molecular dynamics during fibrocartilage differentiation. This data provides a transcriptional resource that will support future investigations of enthesis development at the mechanistic level and may shed light on the strategies for enhanced RC healing outcomes.

## Introduction

Rotator cuff (RC) and its entheses are essential components of shoulder, which are critical in facilitating coordinated shoulder movements and stability[1]. Compared to other RC tissues, the attachment site of supraspinatus (SS) tendon is vulnerable to injury and difficult to achieve complete regeneration, due to its high heterogeneity in composition and structure[1, 2]. Histologically, the attachment site of the SS tendon is a classic fibrocartilaginous enthesis, also termed bone-tendon junction (BTJ)[3, 4]. In its native state, the fibrocartilaginous enthesis exhibits gradations in tissue organization, cell phenotype, and matrix composition[3, 5]. Fibrocartilaginous entheses manage to disperse stress and facilitate load transfer between vastly different materials like tendon and bone, with modulus ranging from 200 MPa to 20 GPa[3]. Unfortunately, the intrinsic regenerative capacity of fibrocartilaginous enthesis is not well-understood, which limits the exploitation of the best and most rigorously proven early intervention programmes[1, 6, 7]. Therefore, understanding the complex process of the enthesis morphogenesis and maturation during development may inform strategies for enhanced BTJ healing.

Currently, the mechanism underlying the growth of the enthesis fibrocartilage is less understood. Details are scarce, but the fibrocartilage layer is formed by a pool of site-specific progenitor cells, and initially organizes as an unmineralized cartilaginous attachment unit[8]. Such development pattern shares an overlapping biologic behavior with the growth plate, which is a process for differentiation of mesenchymal stem cells into chondrogenic cells and then sequentially into fibrocartilage cells[9]. A unique enthesis progenitor pool has been identified since the embryonic stages, as cells sandwiched between primary cartilage and tendon, expressing a mixed transcriptome of both chondrogenic and tenogenic genomic features (Scleraxis and SRY-related transcription factor)[10, 11]. These enthesis progenitors can differentiate into either chondrocytes or tendon fibroblasts, under the regulation of Krüppel-like (KLF) transcription factors[12]. In the later embryonic and postnatal stages, cells from the enthesis progenitor pool ultimately either differentiate into or are replaced by a Hh-positive cell population marked by Gli1 and Ptch1[13, 14]. Gli1^+^ cells and their progenies are retained in the enthesis region throughout postnatal development and eventually populating the entire fibrocartilage region between tendon and bone, thereby contributing to enthesis growth[8, 13, 14]. However, the cell-type composition and cell distribution in the enthesis, as well as biochemical markers for enthesis stem cell/ progenitor used in tissue engineering, remain to be elucidated.

Compared with our in-depth understanding of enthesis development during the embryonic stage, the differentiation of enthesis progenitors during postnatal growth is still not well understood and requires novel methods for elucidation. The normal enthesis maintains a gradient of cell phenotypes,from tendon fibroblast to chondrocyte then to mineralizing chondrocyte and to osteoblast/osteocyte[3, 5, 15]. It is unclear how this gradient in cell phenotypes develops and how it is regulated by the local environment (e.g., ECM, muscle loading, and growth factors)[6]. Single-cell RNA sequencing (scRNA-seq) is a powerful method to identify rare cell populations and deduct the course of differentiation, which can be used to identify various cell types and provide insights into tendon enthesis postnatal development[12, 16]. Here, we applied single-cell transcriptomics to analyze the cellular and molecular dynamics during postnatal tendon enthesis growth. The results provided here for deciphering postnatal tendon enthesis development may facilitate future studies of enthesis regeneration.

## Results

### Neonatal to postnatal day 7 (P7) is the critical stage for enthesis fibrocartilage cell differentiation

The heterogeneity of the fibrochondrocytes inside BTJ has been an open question. Aiming to evaluate the postnatal development of enthesis fibrocartilage, we first stained the shoulder sections with H-E and compared the morphological parameters of the enthesis cells at embryonic day 15 (E15), P1, P3, P7, P14, and P28 (Figure 1a and 1c). At E15, an articular cavity was formed and the supraspinatus tendon (ST) was observed attached to the humeral head. The cells at ST attachment site were highly dense and homogeneous, and were visibly different from tendon cells and the primary cartilage cells inside the humeral head, suggesting that postnatal enthesis is formed by site-specific progenitor cells since embryonic stage. At P1, P3, and P7, the fibrocartilage layer and subchondral bone could hardly be visualized. To examine the cellular morphological changes, we furtherly measured the 2D parameters (including 2D area, frete diameter, and roundness) of enthesis cells, statistics results demonstrated that cell size remarkably increased during postnatal development, from postnatal day 1-7. Notably, the variation of the cell area and diameter also increased during development (Figure 1c). At P14 and P28, the fibrocartilage cells were observed column-like stacked alongside the direction of tendon fiber, with more prominent patterns at P28. Combining all morphological parameters, we performed a principal component analysis (PCA) at different time points (Figure 1d). The first two components appropriately separated the tendon tissues at different time points (e15, 1d, 3d versus 2w and 4w), with 1w located in the middle. Our findings suggested that neonatal to postnatal day 7 (P7) is the critical stage for enthesis fibrocartilage cell differentiation.

**Figure 1.**
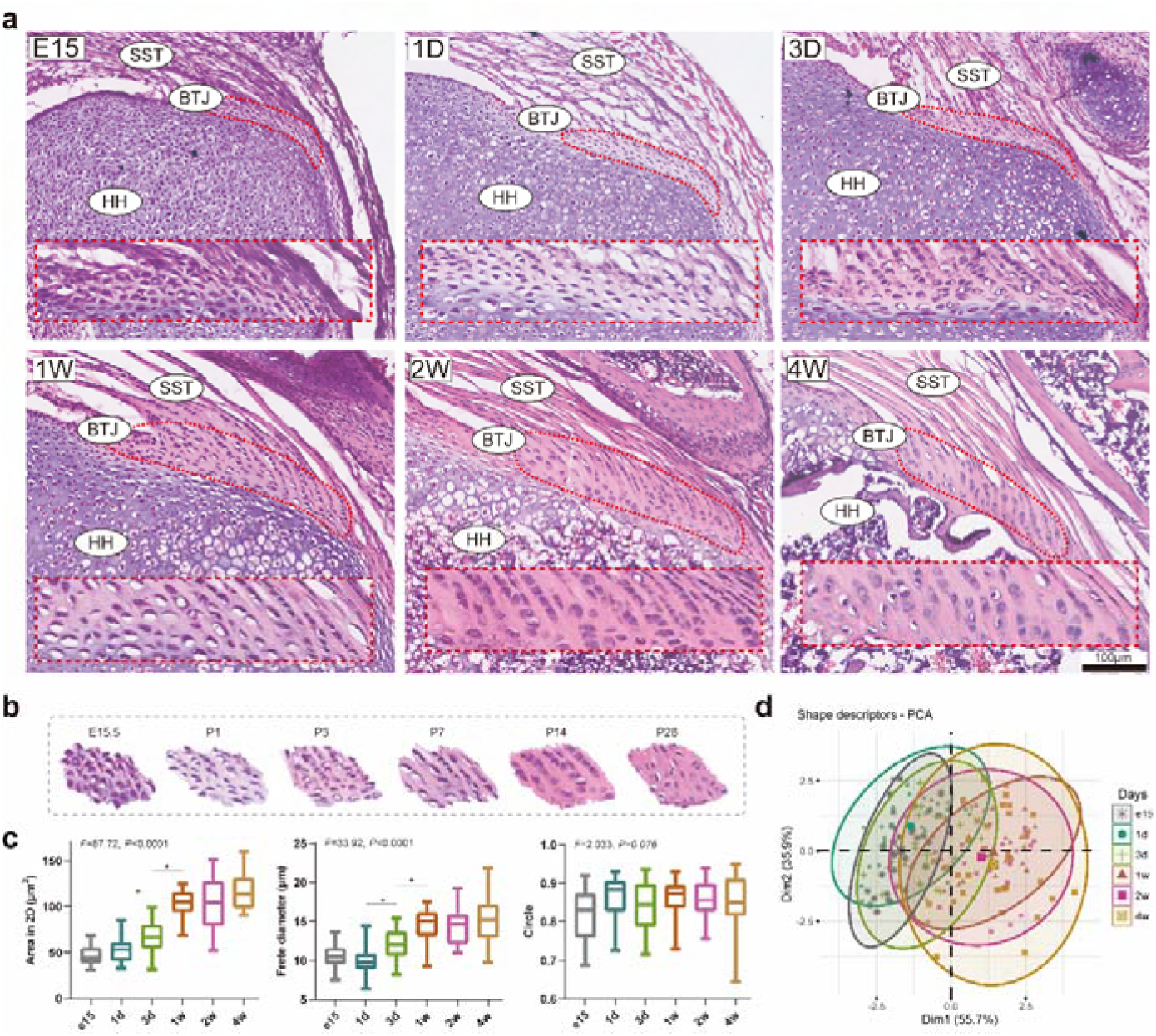
Neonatal to postnatal day 7 (P7) is the critical stage for enthesis fibrocartilage cell differentiation. **(A)** H&E staining of E15, P1, P3, P7, P14, and P28 mouse supraspinatus tendon entheses (n = 4). Scale bars, 100 μm. **(B)** Sketch images of enthesis cells at E15, P1, P3, P7, P14, and P28. **(C)** Comparison of the morphological parameters (2D area, frete diameter, and roundness) between E15, P1, P3, P7, P14, and P28. Error bars represent S.E.M. * = *P* < 0.001 **(D)** PCA of E15, P1, P3, P7, P14 and P28 mouse supraspinatus tendon entheses.

### Unbiased clustering identified known cell populations in postnatal enthesis development

To determine the cellular composition of the developing enthesis, we isolated and sequenced live cells from the developing murine supraspinatus tendon and attached humeral head (Figure 2a). After ruling out the blood cells, endothelial cells, and Cd45+ myeloid cells, we got high-quality transcriptomic data from 15832 single cells, including 5288 E15.5 cells, 4192 P1 cells, 4064 P7 cells, 964 P14 cells, and 1324 P28 cells (Supplementary Figure S1). Unbiased clustering based on Uniform Manifold Approximation and Projection (UMAP) identified 18 major cell populations. Based on the differentially expressed genes (DEGs) (Supplementary figure S2), all the cell clusters were annotated, including three tendon and enthesis-related cell groups: tendon cell (C3), bone-tendon junction cell (C4), and myotendinous junction cell (C5); three articular chondrocyte groups: chondrogenic fibroblast (C8), articular chondrocyte (superficial zone) (C9), articular chondrocyte (intermediate zone) (C10); two growth plate cell groups: proliferating zone/resting zone (C11), hypertrophic zone (C12); three osteogenic cell groups: osteogenic progenitors (C14), osteoblasts (C15), osteocytes (C16) (Figure 2b and s1). The density gradient diagram showed cell identity change from the embryonic to the postnatal stage, indicating that the number of fibroblast-associated cells decreased from the embryonic to the postnatal stage (Figure 2c).

**Figure 2.**
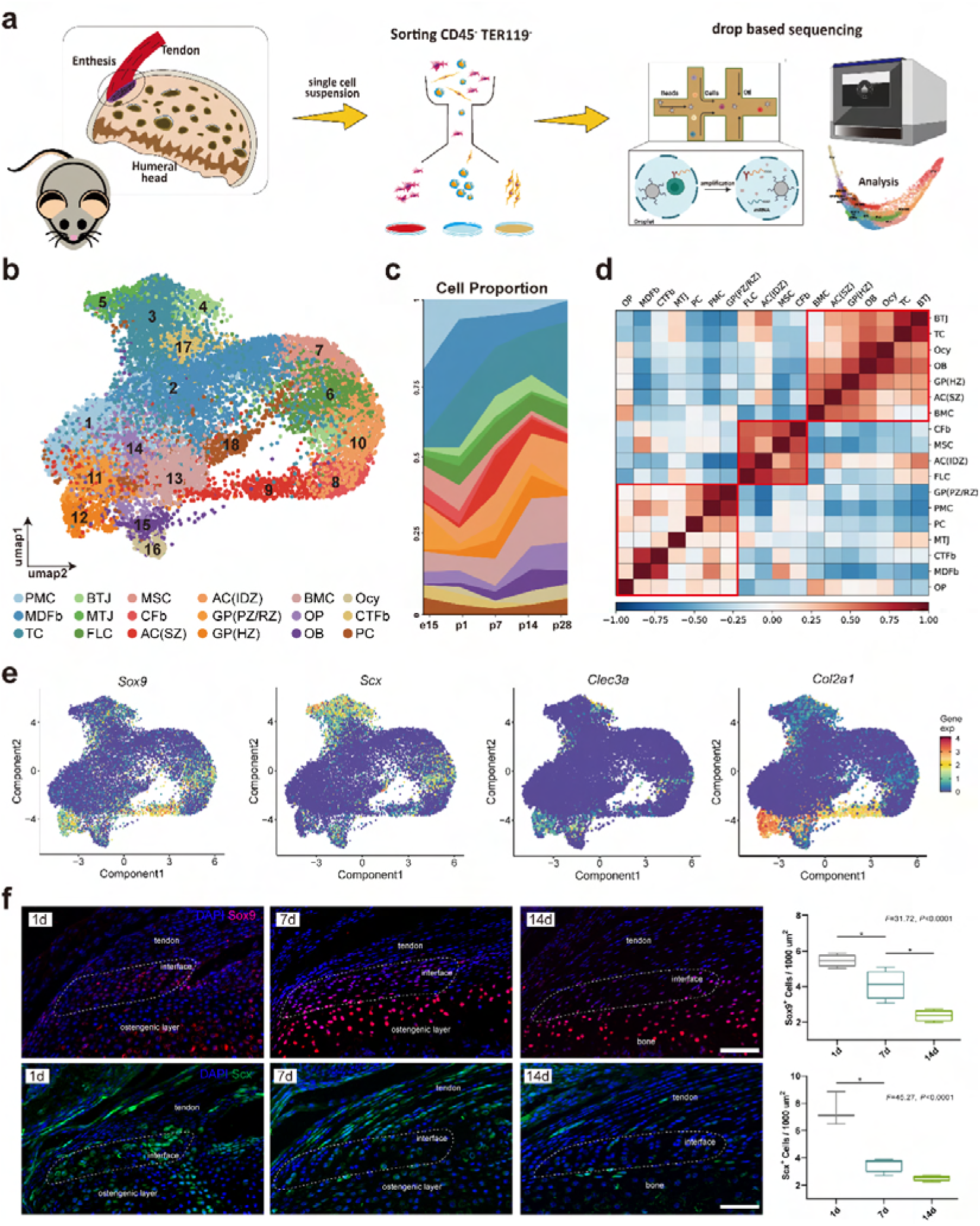
Unbiased clustering identified Known Cell Populations in postnatal enthesis development. **(A)** Schematic workflow of the study design. Cells digested from the supraspinatus tendon, tendon enthesis, and humeral head were subjected to droplet-based scRNA-seq. **(B)** Distributions of 18 cell clusters on a UMAP plot. PMC: primary mesenchymal cell; MDFb: mesenchymal derived fibroblast; TC: tendon cell; BTJ: bone-tendon junction cell; MTJ: myotendinous junction cell; FLC: fibroblast-like cell; MSC: mesenchymal stormal cell; CFb: chondrogenic fibroblast; AC(SZ): articular chondrocyte (superficial zone); AC(IDZ): articular chondrocyte (intermediate zone); GP(PZ/RZ): growth plate cell(proliferating zone/resting zone); GP(HZ): growth plate cell (hypertrophic zone); BMC: bone marrow cell; OP: osteogenic progenitors; OB: osteoblasts; Ocy: osteocytes; CTFb: connective fibroblast; PC: proliferative cell. **(C)** Fraction of cell clusters in enthesis development at E15.5, P1, P7, P14, and P28. **(D)** Heatmap revealing the Pearson correlations between each major cell group in the global transcriptome profile. **(E)** The average expression of curated feature genes for previously reported enthesis marker genes and enthesis-specific ECM genes. **(F)** Representative immunofluorescence staining to validate the spatial distribution of Sox9^+^ and Scx^+^ cells in the enthesis area, at P1, P7, and P14.

Heatmap showed the Pearson correlations between each major cell group in the global transcriptome profile, suggesting that the genome profile of BTJ cells is highly correlative to tendon cells, superficial articular chondrocytes, osteoblasts/osteocytes, and hypertrophic growth plate cells (Figure 2d). We checked the previously reported enthesis marker gene Sox9 and Scx, as well as enthesis-specific ECM genes (Clec3a and Col2a1), which were ubiquitously expressed in bone-tendon junction cell (c4) (Figure 2e). We then performed an immunofluorescence assay to validate the spatial distribution of enthesis-related genes, we found that Sox9^+^ and Scx^+^ cells were detected in the enthesis area, mostly in the neonatal stage and significantly less in postnatal week 2, as expected (Figure 2f).

### Identifying developmental trajectory and regulatory genes during postnatal enthesis growth

We next sought to investigate if the postnatal bone-tendon junction cell development is associated either with tendon development or with primary cartilage development in the humeral head. We first predicted the differentiation state of each cell group from scRNA-seq data by using Cytotrace [16]. The Cytotrace results showed that across all cell clusters, the “stemness” degree of growth plate cells and fibroblasts associated clusters are higher than other cell types (Figure 3a). Among tendon and enthesis-associated cell groups, CytoTRACE scores of the enthesis cells (bone-tendon junction cells) skewed toward a moderate predicted stem potential, which was slightly higher than that in tendon cells, suggestive of a higher degree of stemness for enthesis cells developing into fibrocartilage cells (Figure 3b).

**Figure 3.**
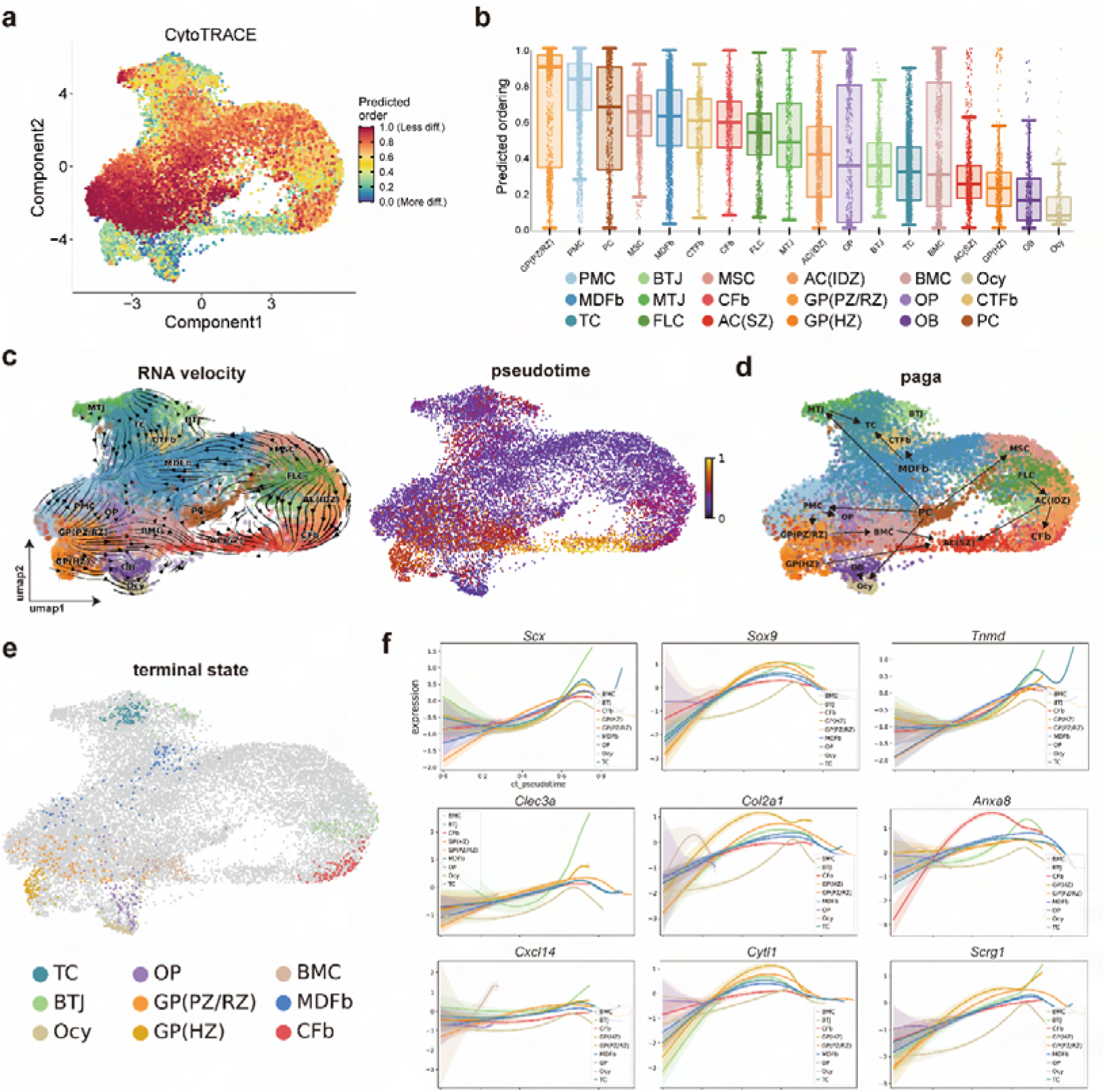
Developmental trajectory and regulatory genes during postnatal enthesis growth. **(A)** UMAP plot of enthesis scRNA-seq data overlaid with CytoTRACE scores. **(B)** Boxplot of predicted differentiation score distributions for each cell cluster. **(C)** Results of RNA velocity analysis show that postnatal enthesis fibrocartilage origin from enthesis site-specific progenitors, instead of tendon cells or the primary cartilage cells that form the humeral head. **(D)** Partition-based graph abstraction (PAGA) shows that the postnatal differentiation of bone-tendon junction cells is independent of tendon cells or other cells. **(E)** Cellrank identified 9 differentiation terminal groups, including bone-tendon junction cells. **(F)** Representative gene expression dynamics along 9 terminal differentiation trajectories.

To resolve the directionality of differentiation between growth plate cells, tendon cells, and bone-tendon junction cells, we implemented RNA velocity analysis. The trajectory results showed four original clusters: growth plate cells, tendon cells, mesenchymal stromal cells, and bone-tendon junction cells (Figure 3c), suggesting that the differentiation profile of these cell groups was separative from each other. Furtherly, the strongest link in directed Partition-based graph abstraction (PAGA) projection further confirms the postnatal differentiation of bone-tendon junction cells is independent of tendon cells or other cells (Figure 3d and Supplementary figure S3). Next, we employed RNA velocity information to compute the terminal states among all the cell groups by Cellrank, which is a toolkit based on Markov state modeling[17]. Cellrank identified 9 differentiation terminal groups, including bone-tendon junction cells. Taken together, the relationship between tendon cells and bone-tendon junction cells, predicted by PAGA and Cellrank, was consistent with RNA velocity, suggesting that the fibrocartilage in postnatal enthesis origin from enthesis site-specific progenitors, instead of tendon cells or the primary cartilage cells that form the humeral head.

To explore gene expression dynamics along the trajectories, we measured the dynamics of genes in pseudotime along the differentiation trajectories of these 9 differentiation terminal clusters (Figure 3f). We found that the expressions of known differential regulator genes were upregulated significantly higher in enthesis-associated trajectories, such as Scx, Sox9, and Tnmd, which were confirmed indispensable for enthesis formation. We also found genes related to ECM organization (Clec3a, Col2a1, and Anxa8) were upregulated along the pseudotime, and highly expressed until terminal differentiation in bone-tendon junction cluster, suggesting the collagen and matrix protein synthesis were predominant in postnatal enthesis growth.

### The functional definition of fibrocartilage subpopulations during postnatal enthesis growth

As enthesis site-specific chondrocytes differentiating into fibrochondrocytes growth is the pivotal step of enthesis development, we sought to determine their cellular heterogeneity, which has been an open question. Our integrated analysis identified 3 subclusters of enthesis cells (BTJ cluster): C1 (chondrogenic fibroblasts), C2 (fibrochondrocytes), and C3 (homeostatic chondrocytes) (Figure 4a). C1 was identified by highly expressed genes (S100a1, Vcan, Postn, Aspn), C2 highly expressed biomineralization-related genes (Clec3a, Tnn, Acan), and C3 expressed Prg4, Col2a1, Ucma, which are markers for articular chondrocytes (Figure 4c). We checked the previously reported enthesis marker genes Col2a1, Col9a1, Clec3a, which were ubiquitously expressed in C2 and C3 (Figure 4d). Subsequently, to track the overall gene landscape changes, nonnegative matrix factorization (NMF) was further applied to identify the molecular classification. All the gene expressions of three BTJ subclusters were divided into 5 metaprograms: Collagen metabolism, Cellular signaling activity, Cellular differentiation, ECM organization, and Cellular proliferation (Figure 4e).

**Figure 4.**
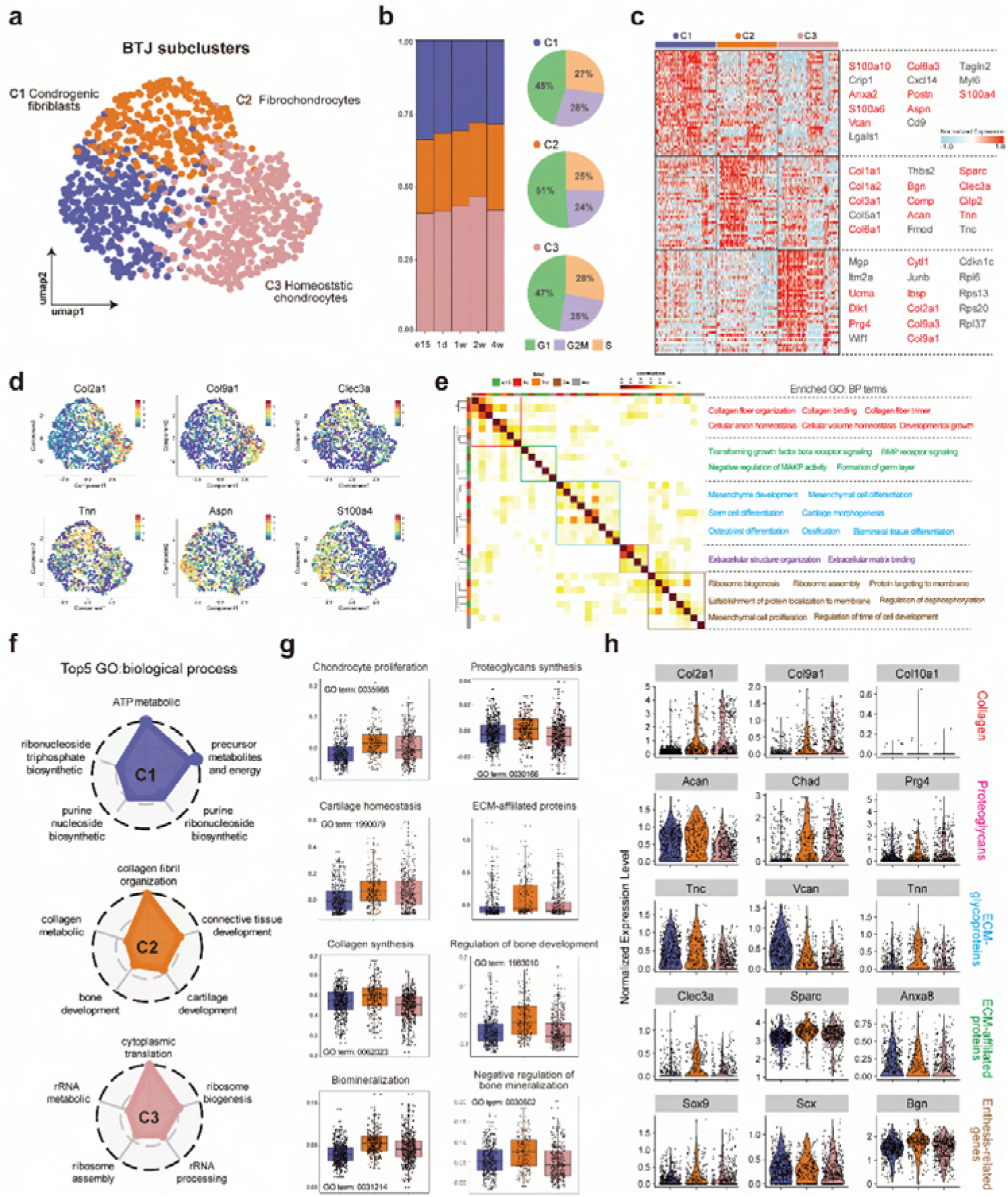
The functional definition of fibrocartilage subpopulations at bone-tendon interface. **(A)** UMAP plot of the three subclusters of chondrocytes in BTJ. **(B)** Fraction of BTJ subclusters at E15.5, P1, P7, P14, and P28 and the fraction of each chondrocyte subcluster arrested in the different cell-cycle phases. **(C)** Heatmap revealing the scaled expression of DEGs for each BTJ subcluster. **(D)** Feature plot showing the curated feature genes for chondrocyte homeostasis, biomineralization, and fibroblastic genes. **(E)** Pearson correlations of expressed metaprogram genes in chondrocytes. **(F)** Radar map showing the top 5 GO annotations of indicated biological process among each chondrocyte subcluster. **(G)** The score of certain biological process-related gene-expressions among each chondrocyte subcluster. **(H)** Violin plots showing the expression levels of representative genes associated with 5 patterns in each chondrocyte subcluster.

To illustrate the biological function of each BTJ subcluster, we next performed Gene Ontology (GO) enrichment analysis. The results showed that C1 dedicated its function to nucleoside metabolism and regulation of the mRNA metabolic process, suggesting its role in postnatal fibrochondrocyte formation and growth. C2 highly expressed cartilage and bone development-related genes, indicating the traits of C2 in collagen synthesis and biomineralization. C3 was annotated for ribosome biogenesis and assembly (Figure 4f). Considering that C3 preferentially expressed ECM-related genes, we chose to classify C3 as homeostatic chondrocytes, which function in maintaining ECM homeostasis. We also evaluated these subclusters using annotated gene sets associated with chondrocyte development and ECM organization (Figure 4g and 4h). The results showed that cartilage ECM-synthesis process (collagen synthesis, ECM-affiliated proteins) exhibited strong enrichment in both C2 and C3. While C2 had the highest score in biomineralization and proteoglycans synthesis.

### Reconstruction of the trajectory and gene dynamics of postnatal fibrocartilage growth

To systematically dissect the fibrocartilage growth process, we then investigated the differentiation potential among the 3 BTJ subclusters based on single-cell transcriptome profiling data. The RNA velocity results were suggestive of C1 being the original cluster, contributing to the differentiation of C2 and C3 (Figure 5a). Together with the Cytotrace results showing that across all BTJ subclusters, cells in cluster C1 were ‘less differentiated’ than the other two clusters (Figure 5b and Supplementary S5a). As UMAP visualization does not maintain the global structure of differentiation dynamics, we next used the Monocle method to determine a pseudotemporal ordering for delineating differentiation paths. The results showing cluster C1 was the start point of pseudotime trajectory (Figure 5c). Inconsistent with the strongest link in directed PAGA projection, pseudotime ordering revealed a transcriptional continuum of C1 cellular states, in which known fibroblastic and proliferative markers towards mature and secretion type (C2 and C3). We hence confirmed that the postnatal formation of BTJ fibrocartilage underwent processes of chondrocyte proliferation, ECM synthesis, mineralization, and cartilage homeostasis.

**Figure 5.**
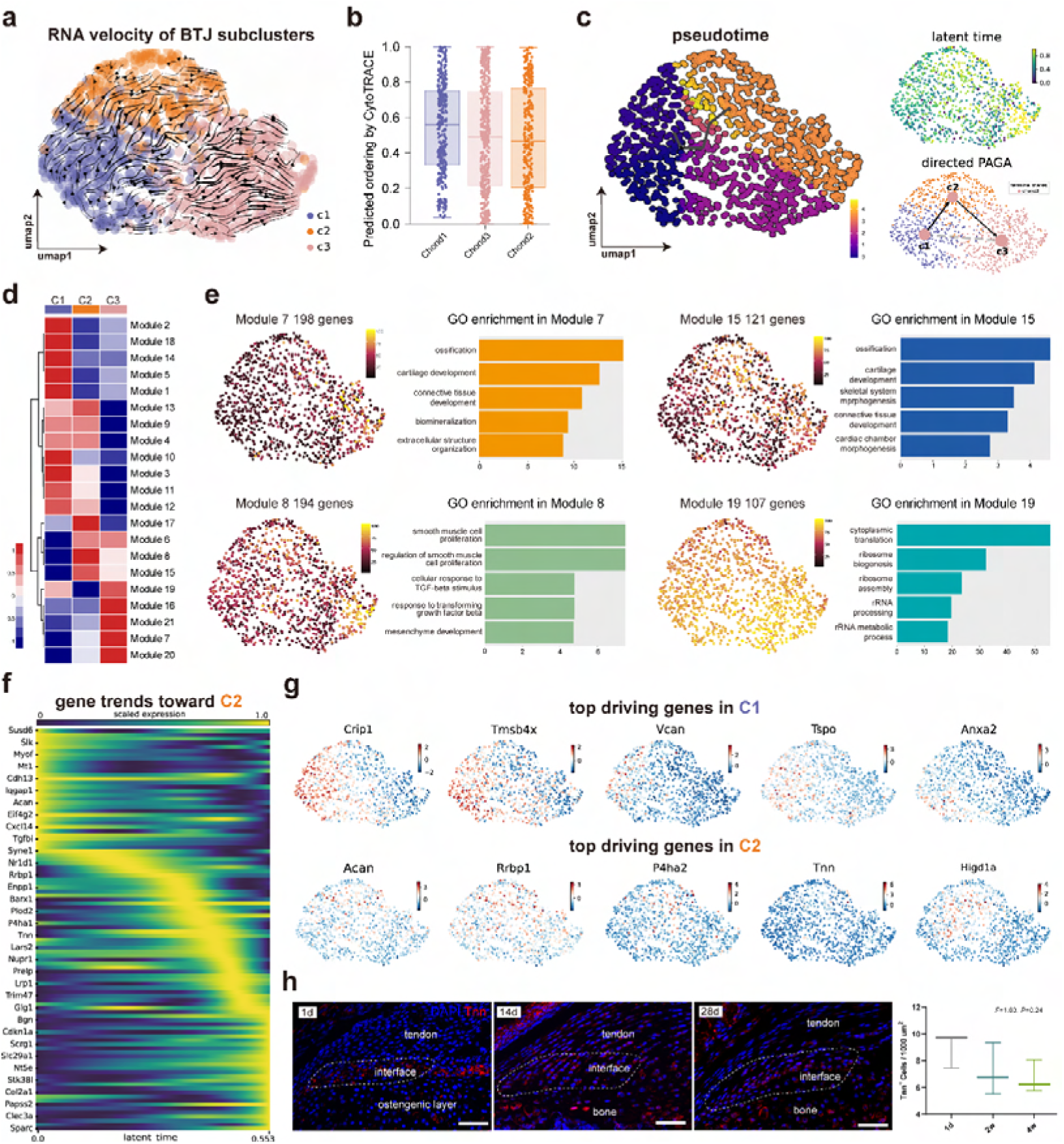
Reconstruction of the trajectory and gene dynamics of postnatal fibrocartilage growth. **(A)** Results of RNA velocity analysis show C1 to be the original cluster, contributing to the differentiation of C2 and C3. **(B)** CytoTRACE scores of the three subclusters of chondrocytes in BTJ. **(C)** Developmental pseudotime of BTJ subclusters inferred by Monocle 3, and PAGA projections showing the postnatal formation of BTJ fibrocartilage underwent chondrocyte proliferation, ECM synthesis, and cartilage homeostasis. **(D)** Heatmap showing the modules of coregulated genes grouped by Louvain community analysis. **(E)** UMAP plots and functional annotation of the representative gene modules among BTJ subclusters. **(F)** Heatmap showing the gene expression dynamics along differentiation trajectories of C2 and C3. **(G)** Featureplots showing the most significant driving genes in C2 and C3 differentiation. **(H)** In-vivo validation of Tnn expression in postnatal BTJ growth.

To explore gene expression dynamics along the trajectories, we next examined gene patterns that varied between BTJ subclusters into 21 modules using Louvain community analysis. We found genes in Module 7 and Module 15 were expressed preferentially in C2 and C3, which were annotated for ossification, cartilage development, and biomineralization. In contrast, chondrogenic genes were less expressed in C1, yet gene modules annotated for cellular proliferation, TGF signaling, mesenchymal development, and ribosome metabolic were preferentially expressed in C1 (Figure 5d and Supplementary S5c). In light of the gene dynamics along the postnatal BTJ fibrocartilage growth, we measured the genes expressed by C2 in pseudotime. The heatmaps show the putative gene programs driving C2 cell differentiation (Figure 5f).

To illustrate the relations between gene expression and different postnatal time stages, we applied the fuzzy c-means algorithm to cluster gene expression profiles of C2 and C3 into time-dependent patterns (from e15.5 to all postnatal stages). In total, we observed 8 distinct clusters of temporal patterns representing genes that are regulated differently (Supplementary Figure S5d). We furtherly determined the driving genes in C2 and C3 differentiation. The results showed that Col9a1, Col11a1, Col5a2, Tnn, and Klf2 were most significant genes in fibrocartilage growth and biomineralization (Figure 5g). According to prior knowledge, except for collagen organization, the expression of Tnn was unlearned for fibrocartilage formation. We next verified the Tenascin N protein in vivo and the results showed that Tnn was specifically expressed in the tendon-bone junction site and located in the fibrocartilage layer (Figure 5h).

### The functional definition of tendon cell subpopulations

Our integrated analysis identified 4 subclusters of tendon cells: TC1 (tendon fibroblasts), TC2 (tendon fibroblasts), TC3 (mature tendon cells), and TC4 (mature tendon cells). TC1 was identified by highly expressed genes (Ly6a, Dlk3, Clec3b), TC2 highly expressed fibroblast-related genes (S100a4, Col3a1), and TC3/TC4 expressed Tnmd, Scx, Mfap4, which are markers for mature tendon cells (Figure 6a and b). The RNA velocity results were suggestive of TC1 being the original cluster, contributing to the differentiation of mature tendon cells (TC3, TC4) (Figure 6c). Subsequently, the overall gene landscape changes of three tendon cell subclusters were divided into 3 metaprograms: Collagen metabolism, Nucleotide and protein biosynthesis, and Myofilament alignment (Figure 6d). The GO enrichment results showed that TC1 dedicated its function to ribonucleoprotein biosynthesis, suggesting its role in cellular proliferation. The ECM-synthesis process (collagen synthesis, ECM-affiliated proteins) exhibited strong enrichment in both C2 and C3 (Figure 6e). Also, to dissect the tendon maturation process, we investigated the differentiation potential among the 4 TC subclusters. The RNA velocity results were suggestive of TC1 was the original cluster, which had the highest differentiation potential, contributing to the differentiation of TC3 and TC4 (Figure 6f and 6h). Furtherly, we measured the genes expressed by TC2 and TC3 in pseudotime. The heatmaps show the putative gene programs driving TC2 and TC3 cell differentiation, respectively (Figure 6h).

**Figure 6.**
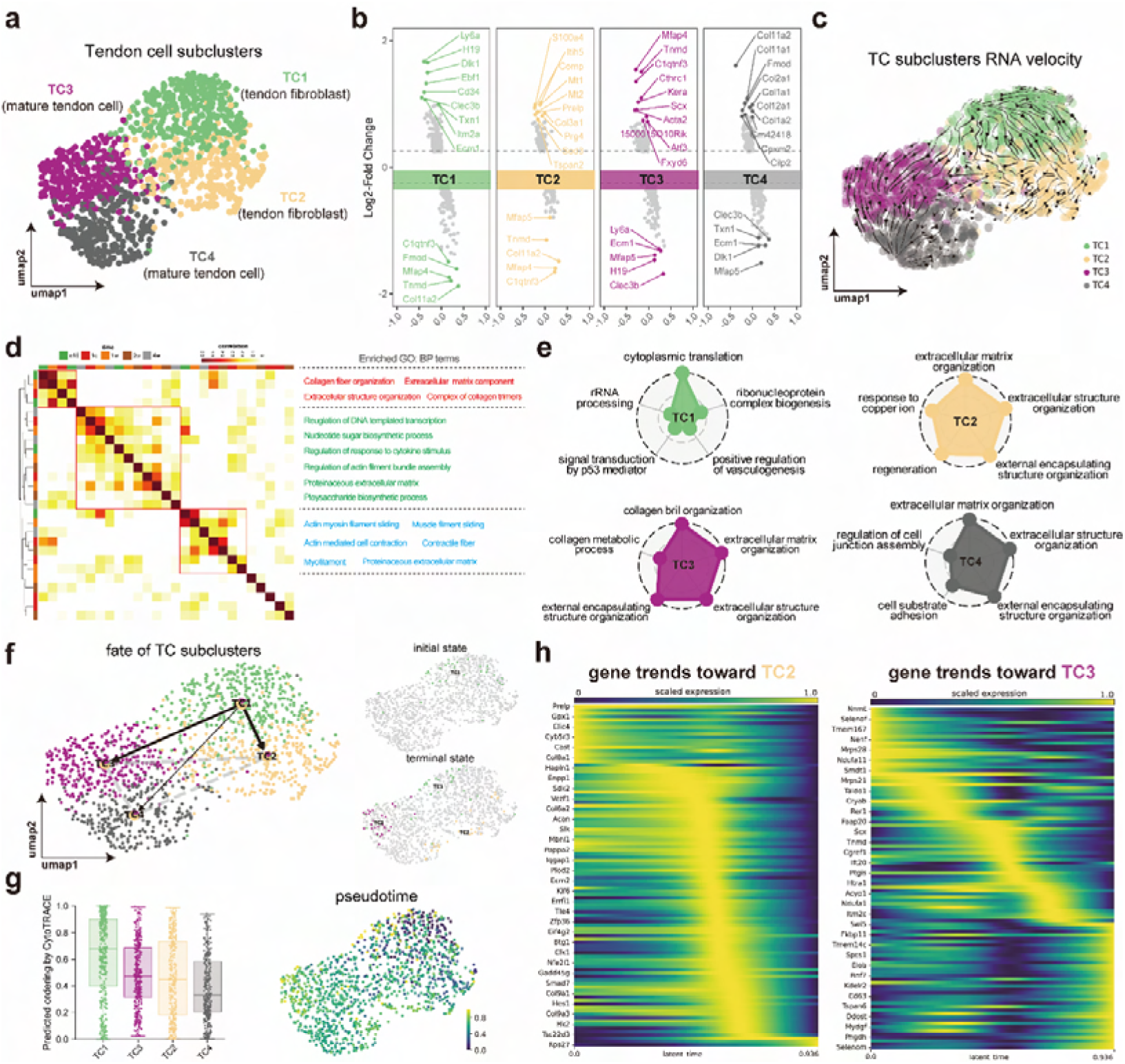
The functional definition of tendon cell subpopulations at bone-tendon interface. **(A)** UMAP plot of the 4 subclusters of tendon cells in BTJ. **(B)** The most upregulated or downregulated genes in each tendon subcluster. **(C)** The RNA velocity results were suggestive of TC1 being the original cluster, contributing to the differentiation of mature tendon cells (TC3, TC4). **(D)** Heatmap showing pairwise Pearson correlations of expressed metaprogram genes in tendon cells. **(E)** Radar maps showing the top 5 GO annotations of indicated biological process among each TC subcluster. **(F)** Partition-based graph abstraction (PAGA) showing the differentiation orientation of TC1. **(G)** Predicted differentiation potentials and pseudotimes of each tendon cell. **(H)** The heatmaps show the putative gene programs driving TC2 and TC3 cell differentiation, respectively.

### Signaling network for the intercellular crosstalk regulating the enthesis postnatal growth

To seek further insights into the critical factors that regulate the enthesis postnatal growth, we performed the signaling network among the 18 subclusters including tendon cells, bone-tendon junction cells, and other cell types. CellChat analysis identified the aggregated signaling network for intercellular crosstalk. Relative active bidirectional signaling interactions among these cell subclusters revealed highly regulated cellular communications (Figure 7a). We then identified the signaling roles of each subcluster, the results showed that bone-tendon junction cells (BTJ cluster) predominately showed incoming patterns, as suggestive of signaling receivers (Figure 7b). We furtherly identified signals that contribute most to the incoming signaling of bone-tendon junction cells (Figure 7c). To determine the important factors, we further analyzed the intercellular signaling networks of FGF and Bmp signaling, which had been reported relative to enthesis development. Firstly, the expression pattern of Fgf-Fgfr signaling was noticed, as both autocrine and paracrine in BTJ cluster. Bone-tendon junction cells, articular chondrocytes, and osteoblasts/osteocytes were leading receivers of Fgf signaling (Figure 7d). We observed Fgfr2 expressed mostly in bone-tendon junction cells and superficial articular chondrocytes via Fgf2-Fgfr2 or Fgf7-Fgfr2. The transforming growth factor-β (TGF-β) superfamily includes a family of proteins, such as TGF-βs (TGF-β1 and TGF-β3) and bone morphogenetic proteins (e.g., Bmp2, Bmp4, and Gdf5). In Bmp signaling network, the bone-tendon junction cells acted as critical mediators and contributors by secreting Bmp ligand Bmpr2, especially (Figure 7e). Specifically, BTJ cluster was the key population that dominated the Gdf5 signaling network.

**Figure 7.**
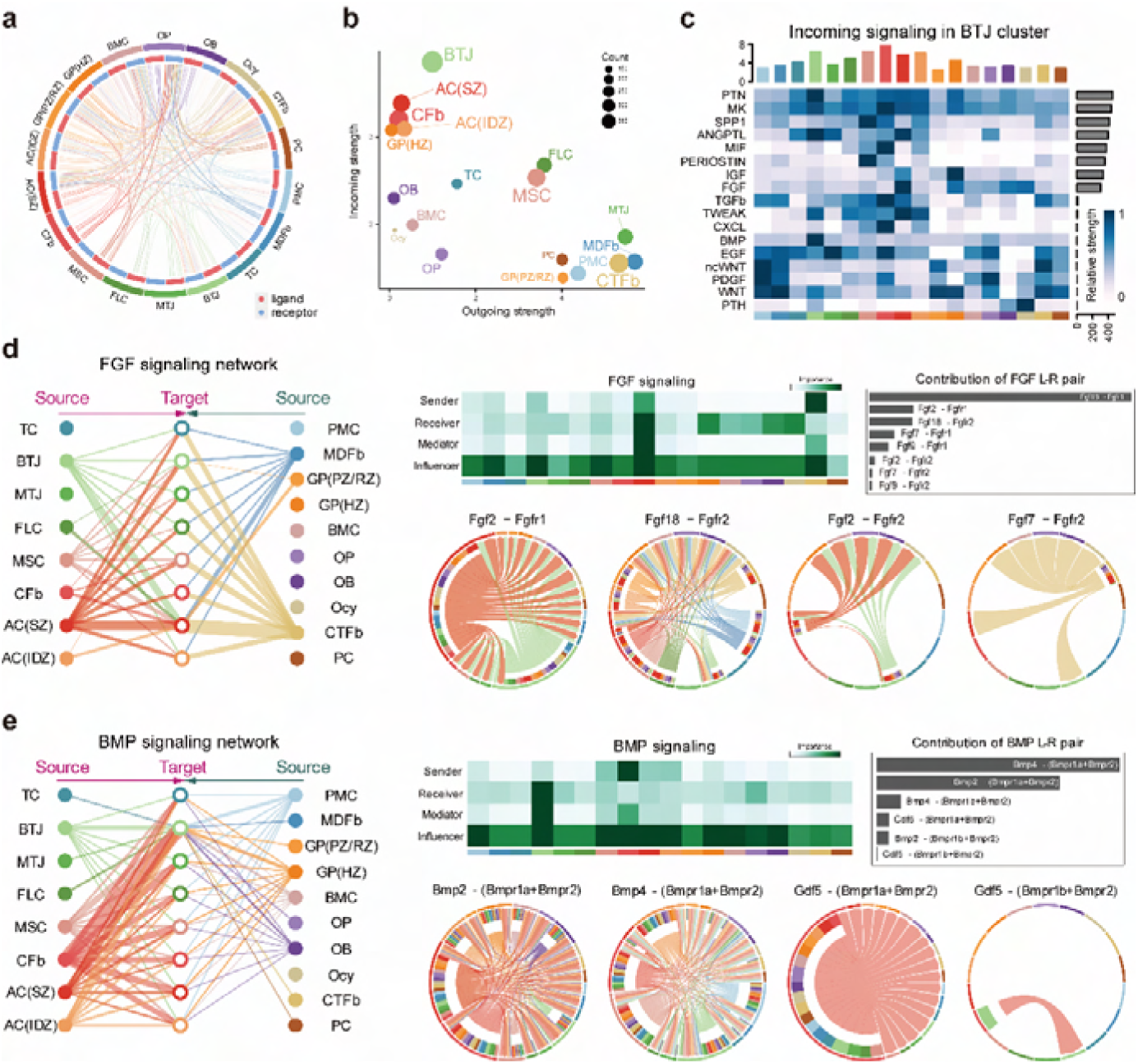
Overview of the crosstalk networks among the clusters. **(A)** Overview of the cellular network regulating the postnatal enthesis growth. Dots indicate cell clusters. The directed line indicates the relative quantity of significant ligand-receptor pairs between any two pairs of cell clusters. **(B)** The dominant senders (sources) and receivers (targets) among 18 cell clusters. **(C)** Identify signals contributing most to incoming signaling of BTJ cluster. **(D)** Chord plots show the inferred FGF (b), BMP (c) signaling networks**. (E, F)** Overview of FGF and BMP signalings networks in enthesis development. Hierarchy plots show the inferred signaling networks among all cell clusters. Heatmaps show the signaling roles of cell groups. Chord plots show the ligand-receptor pairs highly expressed in BTJ cluster.

## Discussion

Deciphering how a complex enthesis is formed from fetal-like into mature status may shed light on the strategies for enhanced BTJ healing[6, 7]. However the cellular complexity and heterogeneity of developing rotator cuff (RC) enthesis are poorly understood, as previous studies could hardly resolve it at the single cell level[18]. So far, there is only one transcriptomic study for embryonic mouse enthesis has been carried out using single cell RNA sequencing[12]. However, the development of fibrocartilage at the enthesis of mouse rotator cuffs occurs no earlier than 2 weeks after birth[19], suggesting the investigation of cellular and genomic mechanisms in postnatal enthesis development is needed. In this study, we applied single-cell transcriptome analysis to delineate the comprehensive postnatal enthesis growth and the temporal atlas from as early as embryonic day 15 up to postnatal day 28. We delineated the cellular heterogeneity and reconstructed the developmental trajectories of postnatal enthesis to describe its functional transformation as well as the underlying molecular cascades. This study may facilitate a better understanding of the enthesis development and add to the single-cell datasets repository of enthesis.

According to prior studies, three distinct populations appear where supraspinatus tendon attaches to the humeral head cartilage: tendon midsubstance progenitors, enthesis progenitors, and primary cartilage progenitors[10, 20]. Enthesis morphogenesis involves predominately enthesis progenitors transforming into fibrocartilage, during which process enthesis progenitors are organized as an unmineralized cartilaginous attachment unit and then mineralizes via endochondral ossification postnatally[19]. Our data showed that developing enthesis cells exhibited a high correlation with tendon cells and chondrocytes from articular surface and growth plate, and the results were inconsistent with the prior studies that postnatal growth of fibrocartilage enthesis consists of cells that express both tenogenic and chondrogenic factors[10, 11]. However, the pseudotime results showed that the directionality of differentiation between tendon cells, articular chondrocytes, and BTJ cells were independent of each other, suggesting that enthesis site-specific progenitors give rise to the fibrocartilage, distinct from tenocytes and epiphyseal chondrocytes. According to the literature, the development of fibrocartilage in the enthesis occurs predominately postnatally, which is not evident at the enthesis of mouse rotator cuffs until 2 weeks after birth[19]. Interestingly, the sizes of enthesis cells were observed remarkably increased during postnatal from postnatal day 1-7, and the principal component analysis revealed that the postnatal day 7 (P7) was the critical stage for fibrocartilage cell differentiation in enthesis. Similarly, Schwartz AG et al reported that Gli1-expressing cells significantly increased and populated at the enthesis site since postnatal day 7, well before the onset of mineralization, and persisted in the mature enthesis[14].

The fibrocartilage cells in tendon enthesis have been traditionally classified into calcified and uncalcified fibrocartilage cells, based on their spatial distribution divided by tidemark[21, 22]. However, the histological classification of the cell population is insufficient, for the reason that the collagen and proteoglycans enriched in fibrocartilage are the key to directional versatile stress reduction, instead of cellular heterogenity[3, 23, 24]. Herein, a deeper understanding of the functional roles of the fibrocartilage subtype is necessary. We divided postnatal BTJ cells into three 3 subclusters based on their GO functions: chondrogenic fibroblasts, fibrochondrocytes, and homeostatic chondrocytes. We first identified chondrogenic fibroblasts, as their highly expressed genes pointed to fibroblast markers (Postn, Aspn, S100a4)[25, 26] and cartilage ECM markers (Vcan, Crip1)[27, 28]. And the gene ontology of chondrogenic fibroblasts was enriched to nucleoside and ribonucleoside synthesis, suggesting their roles were focused on cellular phenotype change. We next identified fibrochondrocytes, as their differential expressed genes were related to chondrocyte-specific ECM proteins (Acan, Comp)[29, 30], as well as genes expressed in fibrochondrocytes (Cilp2, Col1a1, Col3a1)[31–33]. Further, enthesis fibrochondrocytes showed much more enrichment of proteins involved in ECM-affiliated protein synthesis, proteoglycan metabolism, and collagen synthesis. These findings suggested that the main function of enthesis fibrochondrocytes layed in producing molecular components of the fibrocartilage matrix. Similar findings were reported by Deymier AC et al in enthesis proteomic results, which demonstrated the enrichment of proteins involved in proteoglycan metabolism and high ratios of proteoglycans aggrecan, chondroadherin, versican, and biglycan[23]. In contrast, homeostatic chondrocytes showed high similarity to classical chondrocytes (Col2a1, Prg4)[34, 35], which were quiescent, fully differentiated, and responsible for collagen biosynthesis. We also noticed that ribosome biogenesis-related genes were highly expressed in homeostatic chondrocytes, suggesting the synthesis of collagen was very active in these cells.

It is generally recognized that difficulties in restoring the mechanical properties after BTJ injury are largely due to the failure of fibrocartilage recapitulation[36]. Moreover, in modern RC reconstruction surgeries, anchors fix the tendon to the insertion areas without access to bone marrow, which means BMSCs are not likely the main stem cell source for repair[14, 37, 38]. Therefore, it is needed to investigate the native cellular origination and molecular biology of fibrocartilage formation, in order to enlighten developmental engineering strategies. In our scRNA-seq results, chondrogenic fibroblasts were found to exist at the start of the pseudospace trajectory of three BTJ subclusters, which played an important role in the fibrocartilage development, and these chondrogenic fibroblasts were identified since embryonic day 15 in mice. Deepak AK et al isolated mouse embryonic stem cells and proved these cells could differentiate into fibrocartilage chondrocytes, by activation of TGF-β and hedgehog pathways[39]. We found in the trajectory of fibrocartilage development, that chondrogenic fibroblasts initiated with condensation of mesenchymal cells when Vcan is highly transiently expressed. After the induction, differentiating cells synthesize cartilage-specific molecules, including Acan[27]. We also noticed that Tnn was one of the driving genes in chondrogenic fibroblasts differentiating into fibrochondrocytes, and we confirmed the existence of the tenascin N protein (also named tenascin W) in the developing fibrocartilage layer. We still know very little about the basic biology of tenascin W, which have been reported to be expressed in developing and mature bone, specifically in a subset of stem cell niches[40]. Details are scarce, but the stimulating effects of tenascin-W on osteoblastic progenitors’ differentiation and migration have been reported[41, 42], suggesting its potential role in facilitating enthesis progenitors differentiating into fibrochndrocytes.

To investigate the key factors that regulate enthesis development, CellChat analysis was performed. We found that enthesis cells mainly received signals from other cell types, instead of sending signaling factors. And we focused on the growth factors signaling pathways that were involved in the enthesis cells network, mainly including the previously reported FGF family[43, 44], BMP family[10], TGF-β family[7, 45], and PTH family[13, 14] (Figure 7 and Supplementary Figure 6). Among them, we first found that Fgf signaling was widely expressed in enthesis cells. As previously mentioned, the development of fibrocartilage in enthesis was similar to growth plate with an endochondral-like zone. According to the literature, pre-hypertrophic cells in the growth plate express high levels of Fgfr3, and hypertrophic chondrocytes express high levels of Fgfr1[46]. Yet we found enthesis cells mainly expressed Fgfr2, suggesting that the regulation of FGFs in enthesis progenitors’ differentiating into fibrocartilage cells was different from growth plate cartilage. Recent work has confirmed that enthesis development in the mouse mandible is regulated by FGF signaling via Fgfr2-Fgf2 signaling[44]. These findings suggest a potential role of Fgfr2 in enthesis cells differentiating into fibrocartilage during enthesis development. We next noticed the Bmp signaling (specifically Bmp2, Bmp4, and Gdf5) was expressed in enthesis cells, as one key feature of the Sox9- and Scx-positive progenitors is that their dependence on Bmp2 and Bmp4 for specification and differentiation[10, 43]. We found enthesis cells received Bmp ligands predominately from themselves or cells from tendon and articular cartilage. Blitz et al found that Bmp4 derived from the tendon tip induced enthesis progenitors differentiating into chondrocytes, and conditional inactivation of Bmp4 using Scx-Cre blocked formation of the cartilage anlage prefiguring the bone eminence[47]. Interestingly, it has been indicated that BMP-2 could upregulate tenascin-N expression through a p38-dependent signaling pathway[48], and we found Tnn was among the top driving genes in fibrochondrocyte differentiation. Exclusive to Bmp2 and Bmp4, we found enthesis cells received Gdf5 in an autocrine manner. Dyment NA et al employed lineage tracing to confirm that Gdf5 lineage gave rise to fibrocartilage and contributed to the linear growth of the enthesis[20]. We also found the TGF-β signaling in enthesis cells, as TGF-β was important due to its crucial role in cartilage and tendon development[9]. Canonical TGFβ ligands may diffuse to the near tendon and enthesis, facilitating the recruit of chondrogenic cells. Moreover, the secretion of TGFβ1 has been confirmed mechanically mediated[7, 49].

In summary, our study compared the development pattern of enthesis origins with tendon and articular cartilage, then deciphered the cellular complexity and heterogeneity of postnatal enthesis growth and revealed the molecular dynamics during fibrocartilage differentiation, providing a valuable resource for further investigation of fibrocartilage development at the mechanistic level, which may facilitate better understand of the enthesis development and add to the single-cell datasets repository of the enthesis.

## Methods and materials

### Collection of cells from the humeral head-supraspinatus tendon complex

All animal experimental protocols were approved by the Animal Ethics Committee of Central South University (No. 2022020058). The humeral head-supraspinatus tendon samples were dissected from the left shoulders of C57/BL6 mice at embryonic day-15.5, postnatal day-1, day-7, day-14, and day-28. In general, samples were harvested from pooled sibling limbs of two litters (five to six limbs per pool). Following dissection, all the samples were minced immediately and digested in type I collagenase (1mg/ml, Gibico) and type II collagenase (1mg/ml, Gibico) diluted in low-glucose DMEM (Gibco) solution at 37 °C for 30-40 min. Freshly isolated cells were resuspended into FACS buffer containing 2% FBS (Gbico) in PBS. Cell suspensions were stained with antibodies including Ter119-Alexa700 and Cd45-Alexa700 (Biolengend) to remove blood cells. DAPI (BD) stain was used to exclude dead cells. Flow cytometry was performed on BD FACS Aria II, single cells were gated using doublet-discrimination parameters and collected in FACS buffer.

### Droplet-Based scRNA-Seq

8000-10,000 cells were loaded for each age group by Chromium instrument and its chemistry kit V3 (10X Genomics) according to the manufacturer’s guidance. Each cell was encapsulated with a barcoded Gel Bead in a single partition, then amplified to generate single-cell cDNA libraries and sequenced on an Illumina NovaSeq 6000 platform at a sequencing depth of ∼500 million reads. The Cellranger pipeline (version 6.1.1) was used to align the raw reads to the mouse reference genome GRCm38 and to generate feature-barcode matrices. All the low-quality reads were filtered with default parameters.

### Data processing, Quality Control, and Integration

All the feature-barcode matrices were loaded by the Seurat package (v4.1.0)[50], doublets or cells with poor quality were removed (less than 200 genes and greater than 2 Median absolute deviations above the median, or more than 5% genes mapping to the mitochondrial genome). After quality control, all the feature data were scaled with the sctransform algorithm, to avoid unwanted variation including percentages of mitochondrial reads, number of detected genes, and predicted cell cycle phase effect. All the datasets were integrated and batch-corrected by using CCA algorithm, a union of the top 3000 genes with the highest dispersion for each data was set to generate an integrated matrix. Furtherly, this integrated data was analyzed and subclustered to exclude uninterested clusters (including immune cells, red blood cells, endothelial cells, smooth muscle cells, and neural cells).

### Dimensionality Reduction, Clustering, and DEGs analysis

We used the Uniform Manifold Approximation and Projection (UMAP) and Potential of Heat diffusion for Affinity-based Transition Embedding (PHATE)[51] method to visualize the dataset in low dimensions. Furtherly, K-nearest neighbor (KNN) method and the Louvain algorithm were applied to cluster the cells, with 50 PCs selected and resolution set to 2.4, resulting in nine major cell clusters for subsequent analyses. For second-round chondrocyte sub-clustering, we reconstructed the SNN graphs for BTJ clusters, and three subclusters were determined resolution set as 0.6 for each fibrochondrocyte cluster. The FindAllMarkers function in Seurat was used to calculate DEGs among different clusters, the ‘test.use’ function was set to a statistical framework called MAST[52]. Genes met the criteria that 1) expressing in a minimum fraction of 10% in either of the two tested populations; 2) at least a 0.1-fold difference (log-scale) between the two tested populations; 3) adjusted P values less than 0.01, were considered as signature genes. Clusters were annotated according to the expression of those highly variable genes reported in the literature.

### Time-dependent gene signature clustering and NMF clustering

DEGs between different timepoints were acquired FindAllMarkers function in Seurat, temporal pattern analysis and visualization were conducted by using R package Tcseq according to a standard pipeline. After normalizing the above matrix of immune BTJ subclusters, we used the python package pyNMF for Nonnegative Matrix Factorization (NMF) clustering, with rank set to 2:10, the method to brunet, and nrun to 300. Optimal cluster numbers of DEGs were determined based on the cophenetic, dispersion, and silhouette metrics. Through the above algorithm, the time-dependent DEGs were divided into different molecular metagroups.

### Trajectory analysis and cell state analysis

Before trajectory analysis, the S4 Seurat object was transformed into an anndata object using the Seuratdisk package and loaded by Scanpy[53]. Then, all the bam files were processed with Velocyto[54] to quantify the spliced and unspliced mRNA counts. Subsequently, the Velocyto outputs were loaded into scVelo[55] and merged with the anndata object from Scanpy to compute RNA velocity vectors. Low abundance genes (less than 30 total counts) were filtered from the merged dataset. Velocity estimation and PAGA estimation were performed using the “dynamical” mode with default settings. A combined scVelo and PAGA plot were generated using the scvelo.pl.paga() command. The velocity graphs were drawn upon UMAP and PHATE embeddings. After RNA velocity analysis, we used Cellrank[17] package to compute infer the terminal cell state and cluster absorption probabilities using nearest-neighbour relationships and RNA velocity with equal weight in CellRank’s Markov chain model. We set the n_states = 1 when computing the initial states and n_states = 6 when computing the terminal states

### Cell-Cell Interaction analysis

Cell-cell interaction analysis was performed using CellChat[56] package, according to a standard pipeline.

### Gene regulatory network analyses

We applied Single Cell Regulatory Network Inference and Clustering (SCENIC)[57] to identify the cluster-specific gene regulatory networks. The pySCENIC grn method was performed for building the initial co-expression gene regulatory networks (GRN). The regulon data was then analyzed using the RcisTarget package to create TF motifs referring to the mm9-tss-centered-10kb-7 database. The regulon activity scores were calculated using Area Under the Curve (AUC) method. Besides, we used Cellcall package to analyze the cluster-specific TF enrichment and intercellular communication by combining the expression of ligands/receptors and downstream TF activities for certain L-R pairs. Genes that were expressed in less than 10% of the cells of a certain cell type were excluded.

### GO enrichment analysis

GO enrichment of cluster differentially expressed genes was performed using the R package clusterProfiler[58], with a Benjamini–Hochberg (BH) multiple testing adjustment and a false-discovery rate (FDR) cutoff of 0.1. The Gene Ontology Resource database (http://geneontology.org) was used for GO pathway analysis. Module scores for each gene set were calculated using the AddModuleScore function implemented in Seurat. Gene sets used for scoring (Chondrocyte proliferation, Proteoglycans synthesis, Cartilage homeostasis, Collagen synthesis, Regulation of bone development, Biomineralization, Negative regulation of bone mineralization) were selected from the Gene Ontology Browser of MGI Database (C5: biological process gene sets). Visualization was performed using the R package ggplot2.

### Sample harvest and histological observation

The left shoulder of the C57/BL6 mice was harvested at embryonic day 15 and postnatal days 1, 3, 7, 14, and 28. Specimens were obtained and fixed with 10% formalin buffer for 24 hours and rinsed by dual evaporated water then gradually dehydrated by sequential immersion in 70%, 80%, 90%, and 100% alcohol (each for 2 h), finally dried in the air before use. The interface of each SST insertion was cut into small sizes of approximately 1.0 mm * 1.0 mm before high-resolution scanning. After scanning, the samples were embedded in paraffin and then sectioned for histological studies with Safranin-O/Fast green staining. Histologic sections were observed using light microscopy (CX31, Olympus, Germany).

### PCA analysis

PCA analysis of histological results between each time point was conducted by using the R package FactoMineR and Factoextra. The input variables included 4 histological parameters (Cellular area in 2D, Average cellular ferret diameter, Minimum cellular ferret diameter, and Roundness of cells).

### Immunofluorescence staining

The left shoulder of the C57/BL6 mice was harvested at embryonic day 15 and postnatal days 1, 3, 7, 14, and 28. Then fixed with 4% neutral buffered formalin for 24 hours. After decalcifying, dehydrating, and embedded in OCT, specimens were longitudinally sectioned with 10µm. For immunofluorescence staining, the sections were washed with PBS, permeabilized with 0.1% TritonX-100, then blocked with 5% bovine serum albumin (BSA; Sigma-Aldrich, St. Louis, MO). Sections were incubated with primary antibodies anti-Sox9 (Abcam, ab185996, Cambridge, MA), anti-Scx (Abcam, ab58655), and anti-Tnn (Abcam, ab14184) at 4°C overnight, then incubated with Alexa-Fluor 488 conjugated secondary antibody (Abcam, ab150129) and Alexa-Fluor 594(Abcam, ab150120) conjugated secondary antibody at room temperature for 1 hour and counterstained with DAPI (Invitrogen, Carlsbad, USA). All the images were observed and captured using a Zeiss AxioImager.M2 microscope (Zeiss, Solms, Germany) equipped with an Apotome.2 System. Densities of Sox9, Scx, or Tnn positive cells of each captured image were measured using 200x magnification graphs for each slide by the Image J software (National Institutes of Health, Bethesda, MD).

### Statistical analysis

One-way ANOVA and Student’s t-test were performed to assess whether there were statistically significant differences in the results between time groups. Values of *P*< 0.05 were considered to be significantly different. Data were analyzed using Prism 7 software (GraphPad).

## Supplemental Information Titles

Table. S1 Antibodies used in this study.

Table S2 all clusters marker genes, related to Figure 2.

Table S3 BTJ subclusters marker genes, related to Figure 4.

Table S4 BTJ subcluster C2 time-dependent gene cluster, related to Figure S5.

Table S5 NMF_metaprogram_hubgensAndEnrich, related to Figure 4.

Table S6 monocle_module_GO_enrichment, related to Figure 5.

## Acknowledgement

The authors would like to thank professor Hui Xie /Xiang-Hang Luo and other staff from Movement System Injury and Repair Research Center, Xiangya Hospital, Central South University, Changsha, China, for their kind assistance during the experiments.

## Funding

This study was supported by the National Natural Science Foundation of China (NO. 82230085 and 82272572).

## Authorship contributions

Zhang Tao, Wan Liyang: Conceptualization, Methodology, Data curation, Writing - original draft, Writing – review & editing. Xiao Han: Methodology, Data curation, Writing - original draft. Wang Linfeng: Methodology, Data curation. Hu Jianzhong: Writing – review & editing, Project administration. Lu Hongbin: Investigation, Writing – review & editing, Funding acquisition, Project administration.

## Conflict of interests

The authors declare no conflicts of interest relevant to this article.

## Data and code availability

All single-cell datasets created during this study are publicly available at the Gene Expression Omnibus (GSE223751). Any additional information required to re-analyze the data in the paper is available from the corresponding author upon request.

## Abbreviations used

(RC): rotator cuff
(BTJ): bone-tendon junction
(UMAP): Uniform Manifold Approximation and Projection
(NMF): Nonnegative Matrix Factorization
(ECM): Extracellular Matrix

---

**Fig. S1.**
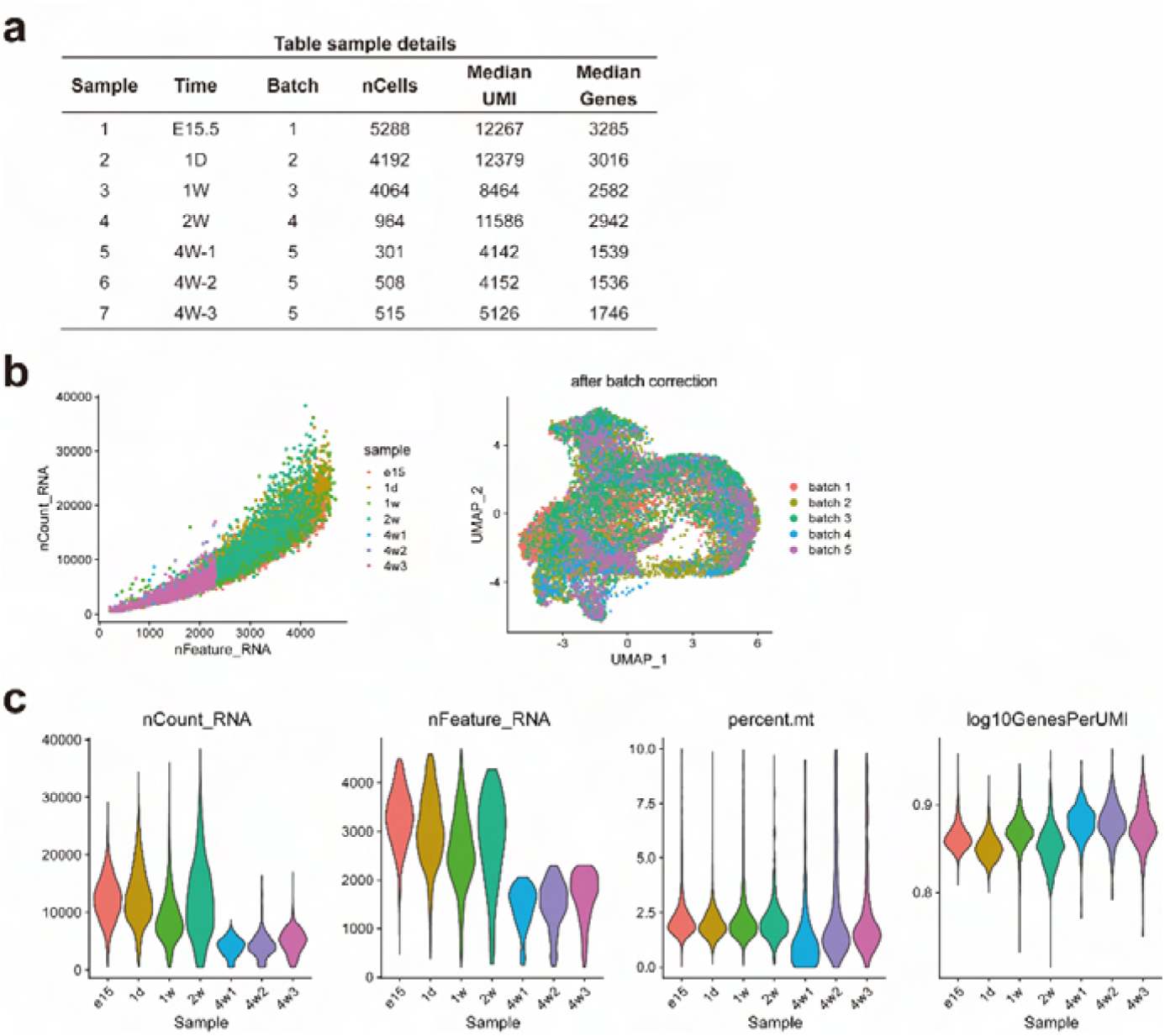
Technical and quality control measures for each scRNA-seq datasets, related to Figure 2. **(A)** Detailed information of each sample. **(B)** Scatter diagram showed the gene numbers (nFeature_RNA) and transcript numbers of single cells (nCount_RNA) per cell after cell filtering; Batch information of single cells from E10.5, E12.5, E15.5 mice hindlimb mapped on UMAP plots. **(C)** Violin plots show the distribution of transcripts and genes detected per cell, percentage of mitochondrial contamination, and sequencing sarutation in each dataset.

**Fig. S2.**
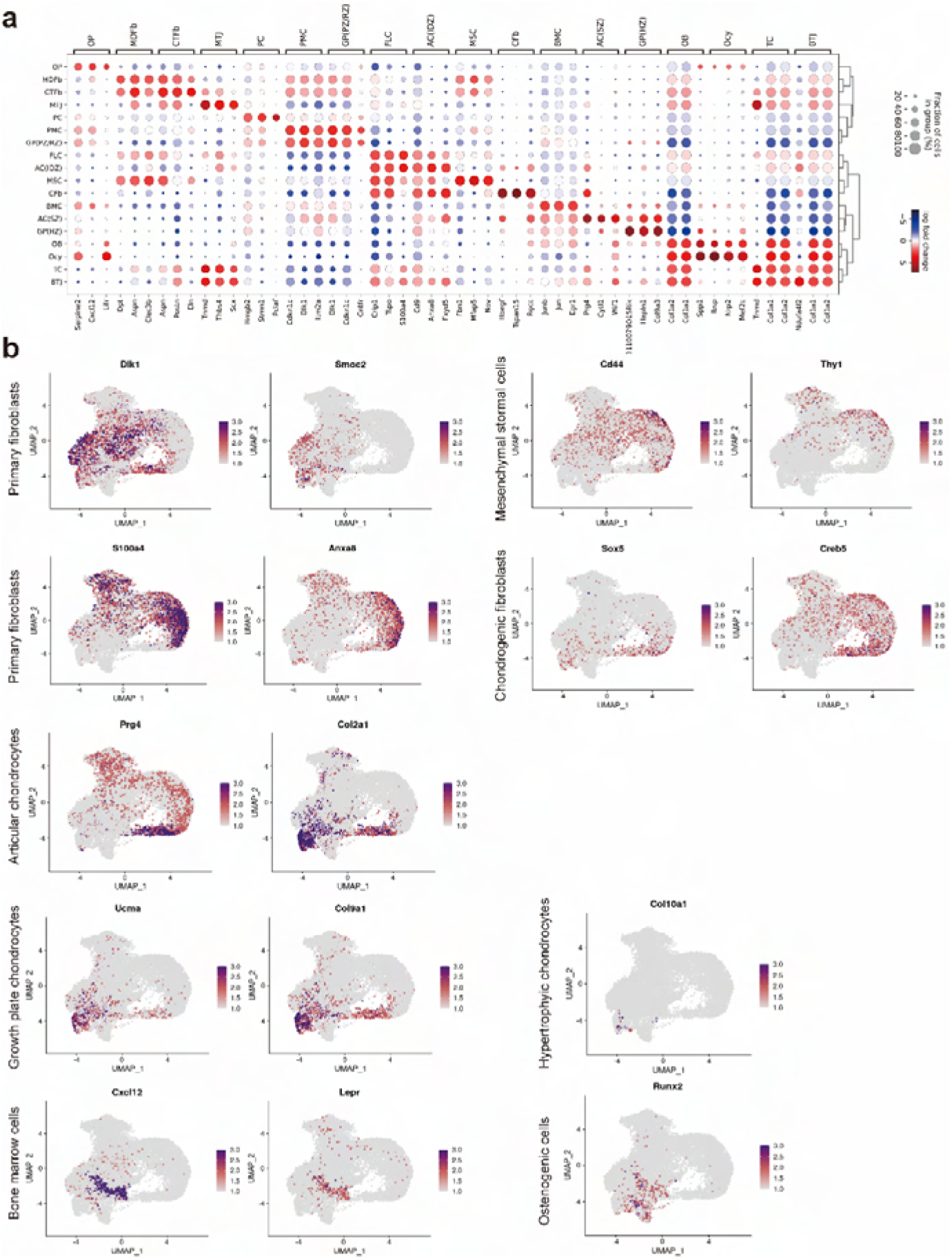
Top differentially expressed genes per cluster in the dataset, related to Figure 2. **(A)** Dotplot of the top 3 highly expressed genes in each cluster. **(B)** The average expression of curated feature genes for previous reported cluster-specific marker genes.

**Fig. S3.**
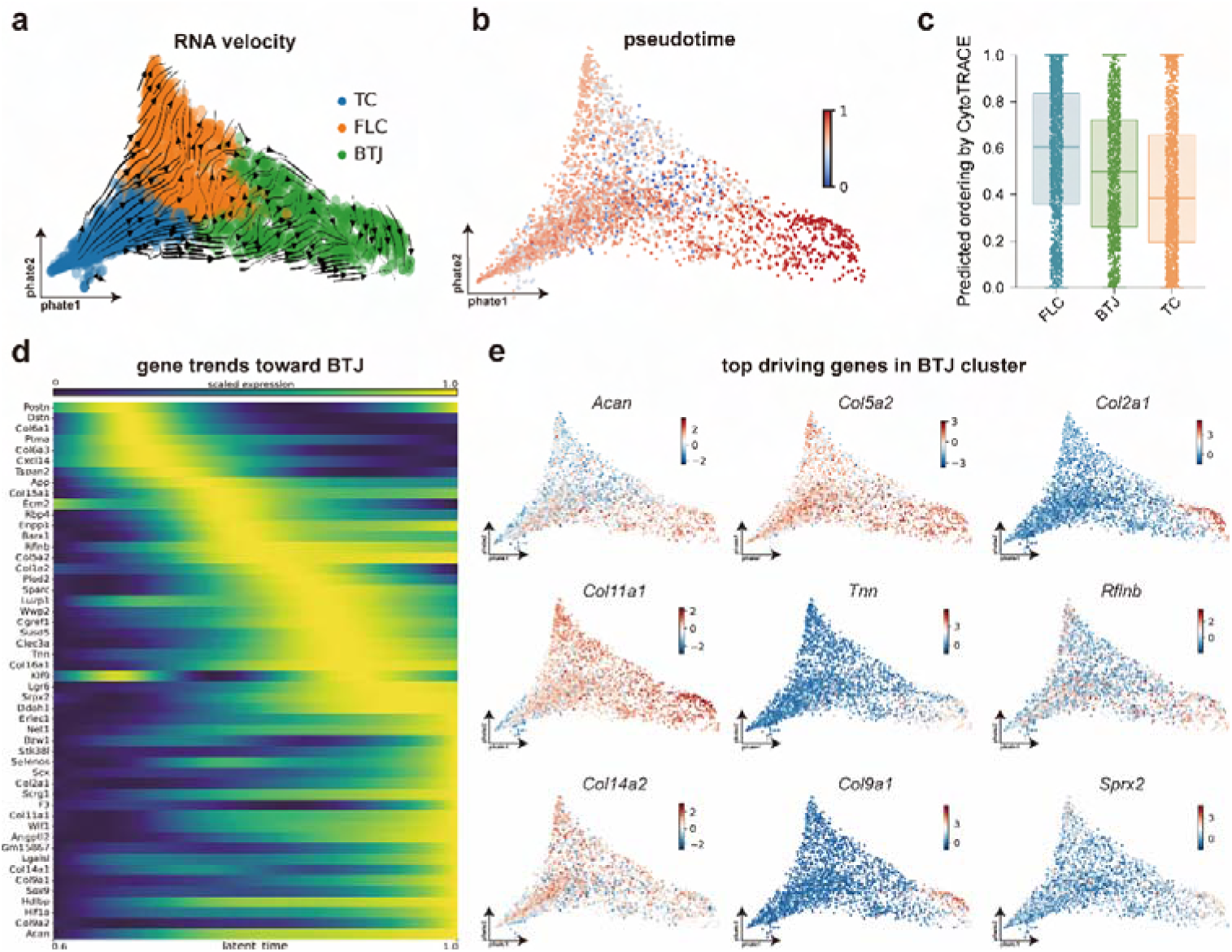
Compare BTJ differentiation with tendon cells, related to Figure 3. **(A)** Results of RNA velocity analysis projected onto PHATE dimensions, showing that postnatal enthesis cells origin from enthesis site-specific progenitors, instead of tendon cells or the primary cartilage cells that form humeral head. **(B)** Predicted differentiation pseudotimes of each clusters’ cells. **(C)** CytoTRACE scores of the three subclusters including enthesis cell. **(D)** The heatmap shows the putative genes that driving BTJ differentiation. **(E)** Featureplots showing the most significant driving genes in BTJ cluster.

**Fig. S4.**
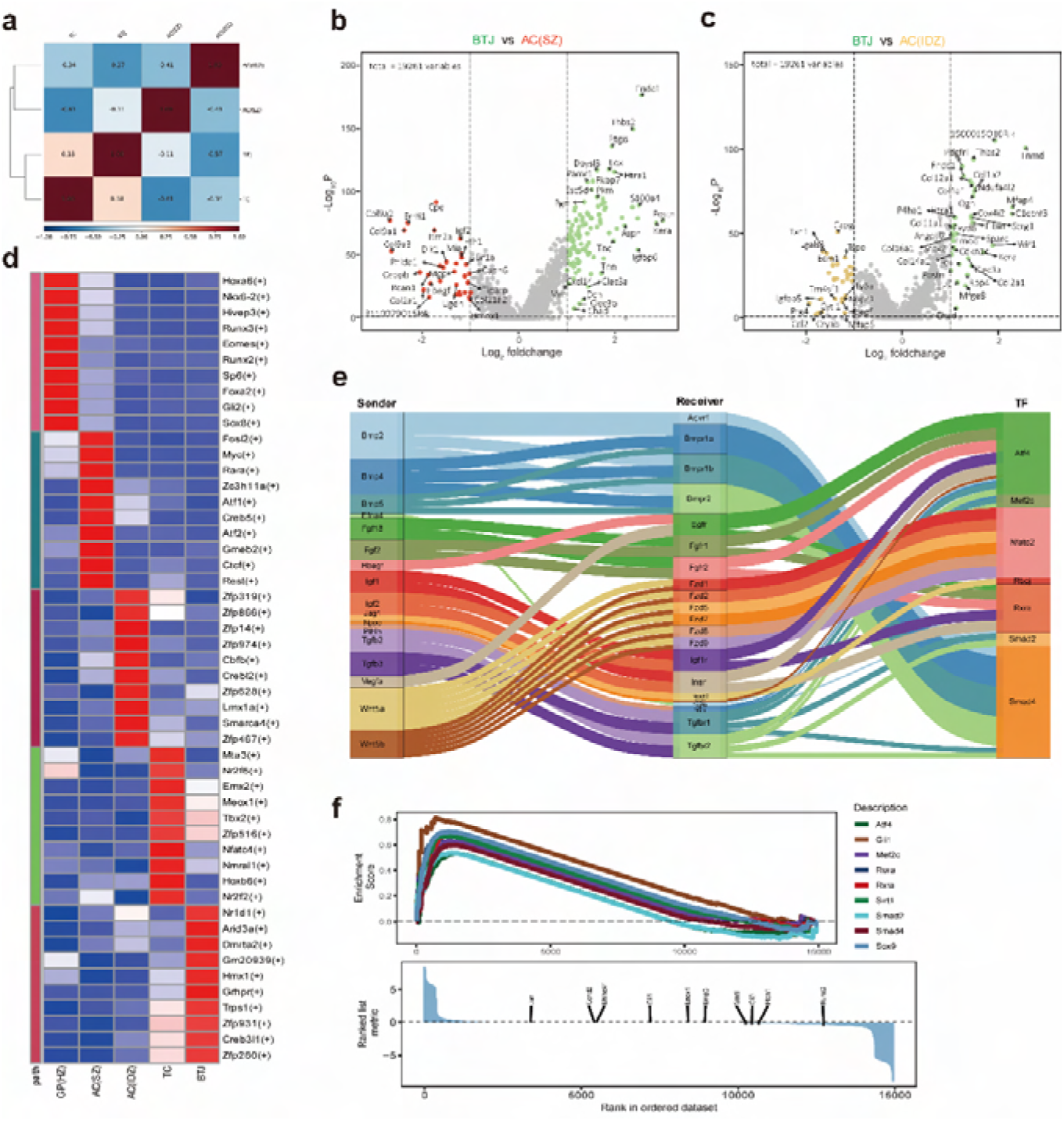
Compare BTJ differentiation with tendon cells and articular cartilage cells, related to Figure 3. **(A)** Correlations between each major cell groups in the global transcriptome profile. **(B, C)** Volcano plots of genes differences between articular chondrocytes and BTJ cells . **(D)** CytoTRACE scores of the three subclusters including enthesis cell. **(D)** Heatmap revealing binary regulon activities analyzed with SCENIC in each subcluster of articular cartilage, tendon and BTJ cells. **(E)** Sankey plot showing the detailed Ligand-Receptor-Transcriptional Factors axis for the communications in BTJ cell types. (F) The top transcriptional factors enriched in BTJ cluster.

**Fig. S5.**
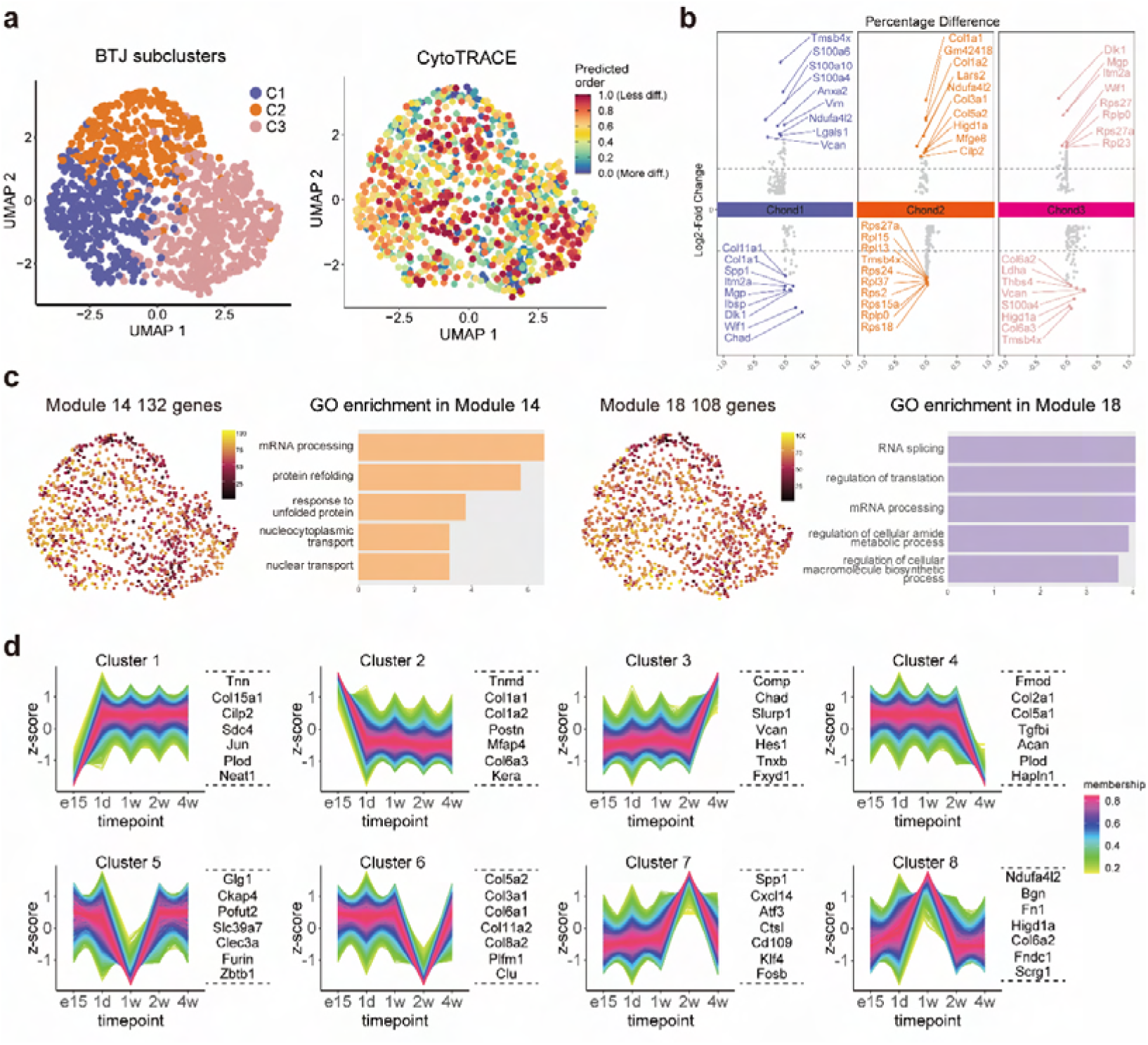
Gene dynamics of postnatal fibrocartilage growth, related to Figure 5. **(A)** UMAP plot of enthesis scRNA-seq data overlaid with CytoTRACE scores. **(B)** The top upregulated or downreguleated genes in each BTJ subcluster. **(C)** UMAP plots and functional annotation of the representative gene modules among BTJ subclusters. **(D)** Fuzzy c-means clustering identified DEGs of C2 into 8 clusters, based on similar expression patterns in differentially expressed timepoint.

**Fig. S6.**
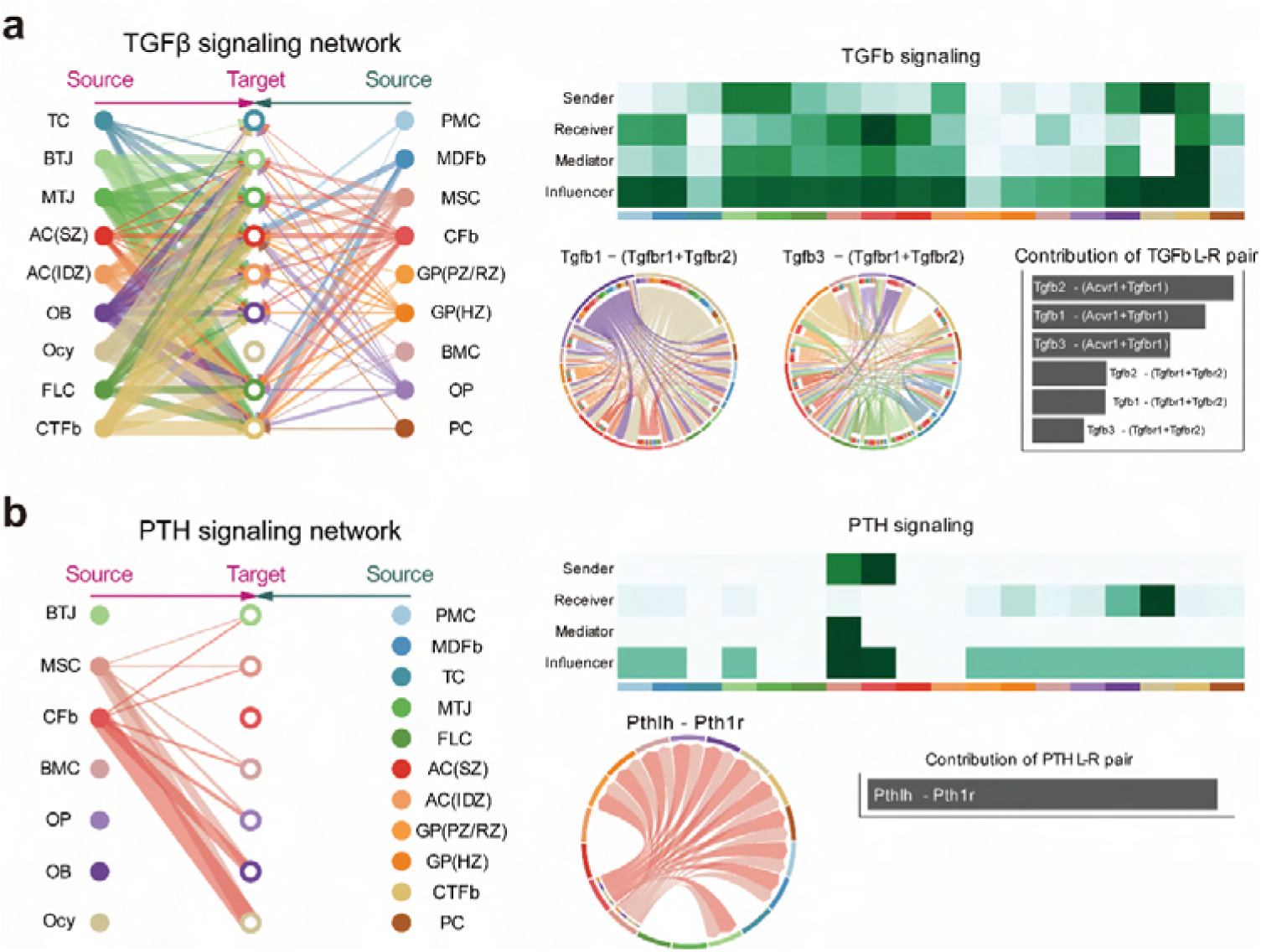
Crosstalk networks of TGFb and PTH signaling among the clusters, related to Figure 7. **(A, B)** Overview of TGFb and PTH signalings networks in enthesis development. Hierarchy plots showing the inferred signaling networks among all cell clusters. Heatmaps showing the signaling roles of cell groups. Chord plots showing the ligand-receptor pairs highly expressed in BTJ cluster.

**Supplementary Table. S1.** Antibodies used in this study.

| **Antibody (clone)** | **Vendor** | **Dilution** | **Usage** |
| --- | --- | --- | --- |
| Alexa Fluor® 700 anti-mouse TER-119 (TER-119) | Biolengend | 0.25ug/million cells | FACS |
| Alexa Fluor® 700 anti-mouse CD45 (I3/2.3) | Biolengend | 0.25ug/million cells | FACS |
| Alexa Fluor® 700 anti-mouse CD31 (390) | Biolengend | 0.25ug/million cells | FACS |
| DAPI | BD | 0.5ul/test | FACS |
| Anti-Mouse Sox9 (ab185966) | Abcam | 1:200 | IF |
| Anti-Mouse Scx (ab58655) | Abcam | 1:200 | IF |
| Anti-Mouse Tnn (DF13237) | Affinity | 1:100 | IF |
| Anti-Rabbit IgG (Alexa Fluor® 488) (ab150073) | Abcam | 1:400 | IF |
| Anti-Rabbit IgG (Alexa Fluor® 594) (ab150064) | Abcam | 1:400 | IF |

## Notes

### Competing Interest Statement

The authors have declared no competing interest.

